# Diaphragm muscle fibrosis involves changes in collagen organization with mechanical implications in Duchenne Muscular Dystrophy

**DOI:** 10.1101/2021.04.07.438870

**Authors:** Ridhi Sahani, C. Hunter Wallace, Brian K. Jones, Silvia S. Blemker

## Abstract

In Duchenne muscular dystrophy (DMD), diaphragm muscle dysfunction results in respiratory insufficiency, a leading cause of death in patients. Increased muscle stiffness occurs with buildup of fibrotic tissue, characterized by excessive accumulation of extracellular matrix (ECM) components such as collagen. However, changes in mechanical properties are not explained by collagen amount alone and we must consider the complex structure and mechanics of fibrotic tissue. The goals of our study were to (1) determine if and how collagen organization changes with the progression of DMD in diaphragm muscle tissue, and (2) predict how collagen organization influences the mechanical properties of ECM. We first visualized collagen structure with scanning electron microscopy (SEM) images and then developed an analysis framework to quantify collagen organization and generate image-based finite-element models. The image analysis revealed significant age- and disease-dependent increases in collagen fiber straightness and alignment, ranging from 4.7 to 13.4%, but collagen fibers retained a transverse orientation relative to muscle fibers. The mechanical models predicted significant age- and disease-dependent increases in transverse effective stiffness and average stress, ranging from 8.8 to 12.4%. Additionally, both healthy and diseased models revealed an increase in transverse stiffness relative to longitudinal stiffness, with significant age- and disease-dependent increases in the ratio of transverse to longitudinal stiffness, ranging from 19.7 to 24.5%. This study revealed changes in diaphragm ECM structure and mechanics during the progression of disease in the *mdx* muscular dystrophy mouse phenotype, highlighting the need to consider the role of collagen organization on diaphragm muscle function.

## 1. INTRODUCTION

### Duchenne muscular dystrophy is a devastating disease with no cure or effective treatment, and diaphragm muscle weakness leads to death

Duchenne muscular dystrophy (DMD) is a fatal genetic disease, with devastating impacts from the subcellular to whole muscle levels. Muscle degeneration results due to a lack in expression of the protein dystrophin, responsible for maintaining the linkage between the intracellular cytoskeleton and extracellular matrix (ECM).^1^ Current therapies are targeted towards either replacing the dystrophin protein or treating the secondary and downstream pathological mechanisms, yet there remains no cure or effective treatment to date.^2^ The absence of dystrophin at the muscle fiber membrane increases susceptibility to mechanical stress from everyday muscle contractions. Regenerative capacity of muscle is decreased, with contraction induced damage leading to a chronic state of inflammation and subsequent fibrosis.^3–7^ A cycle of dysfunction and disuse results in progressive muscle wasting, with differences in severity and progression across muscles.^8^ Lower limb muscle is impacted in the earlier stages of DMD, leading to a loss of ambulation at 10-14 years of age. At later stages of the disease, the diaphragm, the main inspiratory muscle,^9^ is severely impacted. Diaphragm muscle weakness progresses with age in DMD and contributes to respiratory insufficiency, a leading cause of death in the early to mid-20s.^10,11^

### Fibrosis limits muscle function, but the structure and mechanics of fibrotic tissue are not well characterized in diaphragm muscle

One of the primary sources for progressive muscle dysfunction in DMD is the development of fibrosis, characterized by excessive accumulation of extracellular matrix (ECM) components such as collagen. Indeed, many studies have shown that dystrophic muscles have increased amounts of collagen,^12,13^ but collagen amount does not correlate with tissue stiffness in diaphragm muscle tissue from *mdx* mice, the most common animal model used to study DMD.^14^ *Mdx* lower limb muscle shows an increase in collagen fiber alignment.^15^ While collagen fiber amount does not predict tissue stiffness, collagen fiber alignment is reported to be a significant predictor of passive lower limb muscle stiffness.^15^ However, the *mdx* lower limb muscle does not mimic the severity of the human phenotype nearly as well as the diaphragm does. Similar to the human condition, *mdx* mice exhibit impairment in respiratory function^16,17^, decrease in diaphragm muscle fiber cross-sectional area, and increased diaphragm muscle fibrosis.^18–21^ Beyond skeletal muscle, changes in collagen organization with fibrosis are implicated in additional tissue systems. Increased collagen fiber alignment is reported in pulmonary fibrosis^22^ and increased collagen fiber straightness is reported in cancerous pancreatic tissue.^23^ Changes in collagen organization, such as collagen fiber direction, alignment, and straightness, remain unknown in *mdx* diaphragm muscle, but are needed to understand how and why diaphragm muscle mechanics change during the progression of DMD.

### Finite element models allow us to study structure-function relationships in biological tissues

In prior studies, collagen organization is related to passive properties of skeletal muscle measured by mechanical testing of intact muscle.^14,15^ The contribution of the ECM to the passive properties of skeletal muscle is shown by indirect measurements, comparing properties of single muscle fibers and muscle fiber bundles with intact ECM,^24–26^ and direct measurements of properties of decellularized muscle fiber bundles.^27^ While it is difficult to isolate the influence of organizational parameters on ECM properties with traditional mechanical testing, finite-element (FE) models allow us to isolate the impact of specific structural variations on mechanical properties. Whole muscle-level FE models reveal the influence of macroscopic muscle architecture on mechanical properties^28^ but have been limited in their representation of the ECM. These models typically lump together connective tissue, muscle fibers, and muscle fascicles into one transversely isotropic material, without accounting for changes in ECM structure during fibrosis.^28–31^ Muscle fascicle-level FE models reveal transversely anisotropic behavior^32^ and when changes in microstructure due to DMD were simulated, their influence on macroscopic properties was dependent on the relative stiffness between the ECM and muscle fibers.^33^ However, these muscle fascicle-level FE models assumed that the ECM was aligned with muscle fibers and did not account for changes in the structure of the ECM during DMD. Therefore, as we apply these modeling techniques to study fibrotic muscle, we must first consider the complex structure and function of the ECM.

### We aim to quantify changes in ECM structure and mechanics during the progression of diaphragm muscle fibrosis

The goals of our study were to (1) determine if and how collagen organization changes with the progression of DMD in diaphragm muscle tissue, and (2) predict how collagen organization influences the mechanical properties of ECM. We aimed to characterize collagen organization within the epimysium and predict how changes in its structure are implicated on both transverse (cross muscle fiber) and longitudinal (along muscle fiber) tissue properties during disease progression. To do so, we developed an image-based finite-element modeling pipeline to explore the influence of collagen organization on ECM mechanics. We collected scanning electron microscopy (SEM) images of epimysium isolated from *mdx* and wild-type (WT) control mice at 3, 6, and 12 months, and quantified collagen fiber direction, alignment, and straightness. We then generated finite-element models and simulated biaxial stretch to determine the implications of our collagen organization measurements on ECM mechanical properties.

## 2. METHODS

### Animal protocol

All experiments were approved by the University of Virginia Animal Care and Use Committee. This study was conducted in C57BL/10ScSn-Dmdmdx/J male mice (referred to as *mdx*), bred inhouse, and C57BL/6J male mice (referred to as WT), purchased from Jackson Laboratories. Our study groups included 3, 6, and 12-month-old mice, both WT and *mdx* (n=6 mice per group).

### Ex vivo sample collection and imaging

After humane euthanasia, the diaphragm muscle was excised and samples from the costal region were dissected and placed in phosphate-buffered saline. We followed a standard sodium hydroxide digestion protocol to digest muscle fibers and leave only collagen fibrils, removing the sarcolemma, basement membrane, and proteins associated with the ECM.^34^ Excised muscle samples were first placed in fixative for 24 hours (8% glutaraldehyde,16% paraformaldehyde, 0.2M sodium cacodylate). Samples were then placed in digestion solution (10% sodium hydroxide) for approximately 6 days, and then rinsed in H_2_O for 24 hours. Since samples were physically unconstrained during digestion, this protocol left the tissue in a zero-strain configuration after muscle fibers were digested. Therefore, we assume that the arrangement of collagen fibers reflects their position at muscle fiber rest length. Samples were prepared for Scanning Electron Microscopy (SEM) imaging following standard dehydration with a graded series of EtOH (10-100%), mounted on 1/8” stubs, and sputter coated in gold. Care was taken to ensure that tissue samples were mounted such that SEM images of the surface plane captured the outer epimysial layer of the ECM. One image per sample was captured at the following magnifications: 40X, 500X, 1kX, 15kX to visualize collagen organization (Zeiss Sigma VP HD field SEM). Images collected at 1kX were then used for measurements in our image analysis. SEM images collected in this study will be made publicly available upon publication.

### Image analysis

#### Muscle fiber direction

After digestion in sodium hydroxide, collagen structure was isolated from samples of diaphragm muscle. Ridges in tissue samples were still evident, indicating the presence of muscle fibers that had been digested from the samples. From these features, we measured muscle fiber direction manually from raw images (1024×768 pixels) by tracing three locations along the length of digested muscle fibers and averaging the angles of the traces (**Fig. 1A**). Images collected at 1kX magnification were rotated such that the muscle fiber direction aligned with the horizontal direction, and then the images were cropped to a square (540×540 pixels) (Adobe Illustrator) (**Fig.1B**). Images collected at 15kX were used to confirm successful digestion by visually inspecting that muscle fibers were not present (**Fig. 1C**).

**Figure 1.**
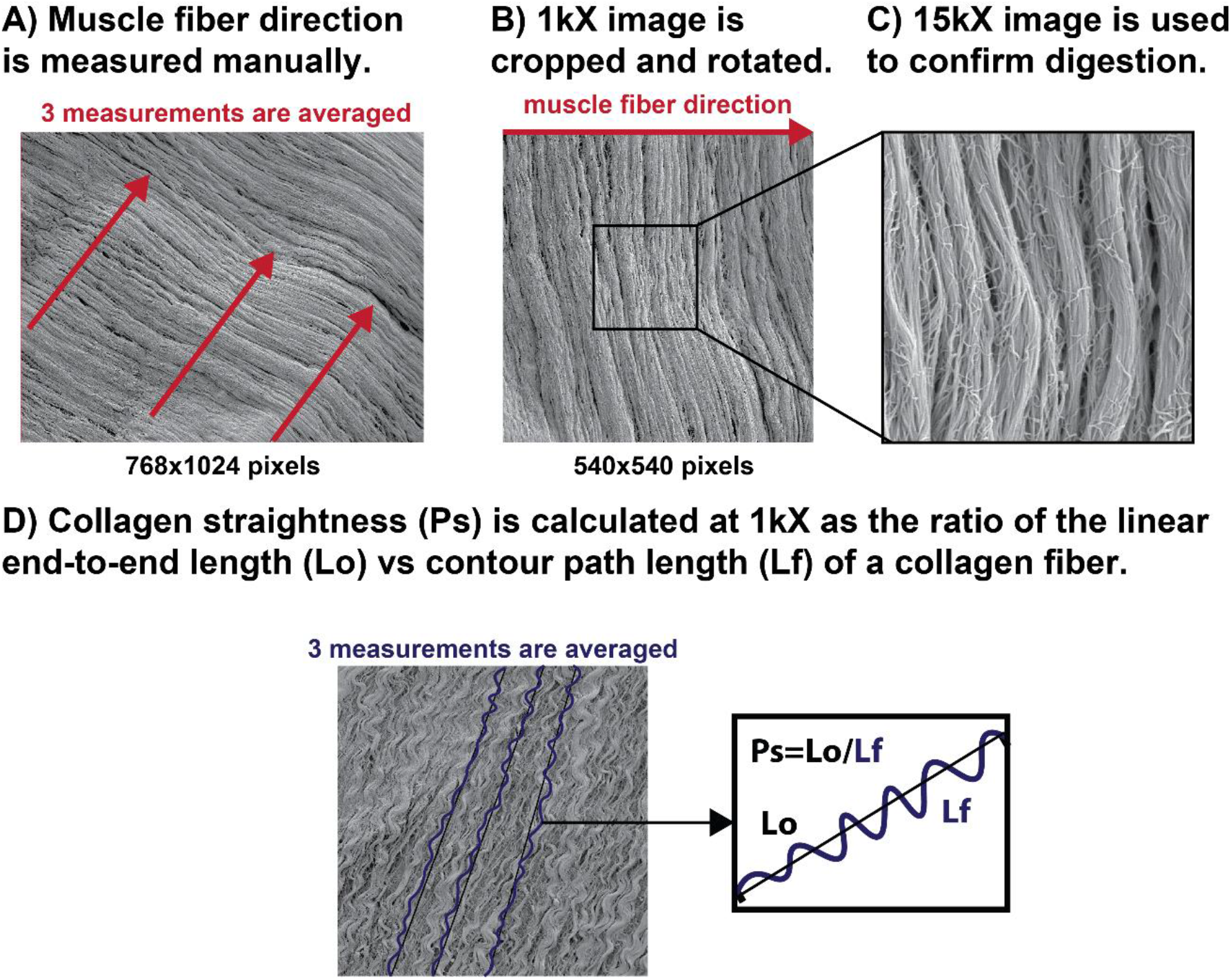
(A) After digestion in sodium hydroxide, collagen structure was isolated from samples of diaphragm muscle. Ridges in tissue samples were still evident, indicating the presence of muscle fibers that had been digested from the samples. From these features we measured muscle fiber direction manually from raw images (1024×768 pixels) by tracing three locations along the length of digested muscle fibers and averaging the angles of the traces. (B) Images collected at 1kX magnification were then rotated with muscle fiber direction on the horizontal and cropped to a square (540×540 pixels) (Adobe Illustrator). (C) Images collected at 15kX were used to confirm successful digestion by visually inspecting that muscle fibers were not present. (D) Collagen straightness parameter (*Ps*) was determined by manually tracing the contour path length of a representative collagen fiber (*Lf*) and the linear end-to-end straight-line length was determined by drawing a straight line connecting the ends of the measured fiber (*Lo*). Collagen straightness parameter (*Ps*=*Lo/Lf*) was then calculated for three fibers representative of the fibers in each image and averaged (ImageJ).

#### Collagen fiber straightness

Collagen straightness (*P_s_*) was determined using the following relationship:

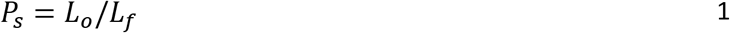

where *L_f_*, collagen fiber length, was calculated by manually tracing the contour path length of a representative collagen fiber, and *L_o_*, the linear end-to-end straight-line length, was determined by drawing a straight line connecting the ends of the measured fiber. The collagen straightness parameter was then calculated for three representative fibers and averaged (ImageJ) (**Fig. 1D**).

### Image processing algorithm

#### Local collagen measurements

We developed an image processing algorithm to automatically measure collagen orientation within subregions of each image in MATLAB (MathWorks Inc., Natick, MA). Each image was discretized into 4,096 sub-regions of 16×16 pixels (**Fig. 2A**). We utilized built-in image processing functions (MATLAB R2018b and Image Processing Toolbox 3.5.8) to measure collagen fiber direction. Each sub-region of the image (*i*) was first thresholded using Otsu’s method.^35^ Collagen pixel ratio (*cpr_i_*) was calculated by dividing the number of white pixels detected as collagen fibers (*n_coll_*) by the number of total pixels (*n_tot_*) (**Fig. 2B**).

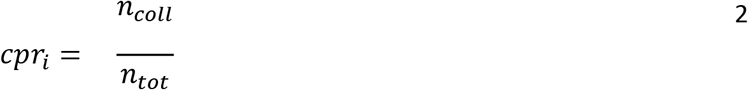

**Figure 2.**
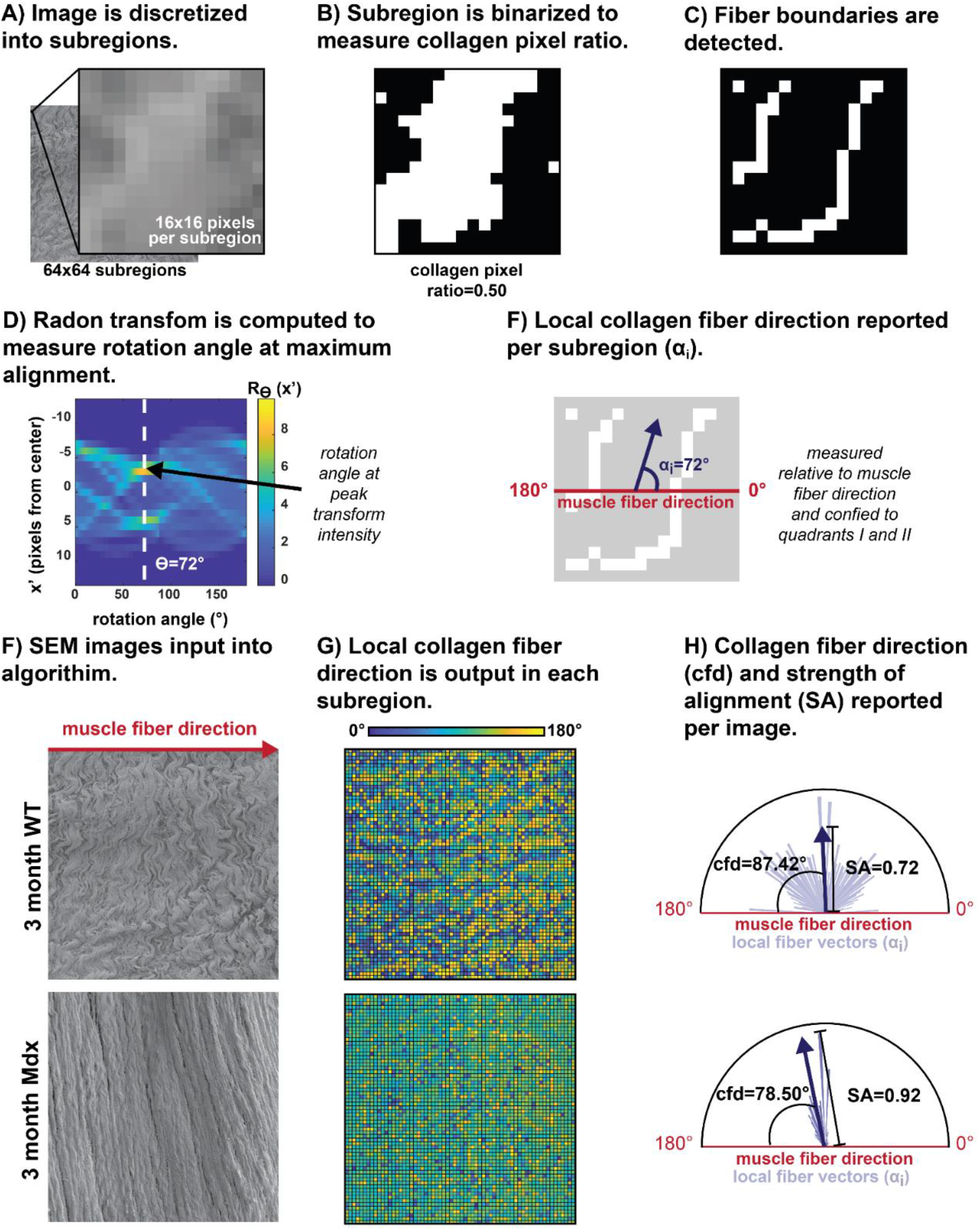
(A) Scanning electron microscopy images were discretized into 64×64 image subregions (16×16 pixels per subregion). (B) Each subregion was thresholded (Otsu’s Method) and collagen pixel ratio was measured (*n_coll_/n_tot_*). (C) Fiber boundaries (Canny Edge Detection) were determined. (D) The Radon transform was computed at fiber boundaries, with rotation angle at the maximum peak used as a measure of dominant orientation. (E) Local collagen direction (*α_i_*) was reported per image subregion relative to the horizontal axis (muscle fiber direction) and constrained to the first two quadrants. (F) Examples of input images for 3-month-old WT (used above in E-F) and *mdx* mice. (G) Local collagen directions measured per image window displayed with a spatial heat map for each image. (H) Mean collagen fiber direction (*cfd*) was reported per image by taking the circular mean of local collagen directions (*α_i_*), measured as the acute angle relative to the horizontal axis (muscle fiber direction). Strength of alignment (*SA*) was quantified as the length of the mean resultant vector and used to measure the circular spread in local orientations per image (*0*<*SA*<*1*, 1=high alignment).

Fiber boundaries were determined from thresholded image subregions using canny edge detection (**Fig. 2C**). We then computed the Radon transform^36,37^ at fiber boundaries to measure fiber orientation^38–40^. The Radon transform, *R_ϴ_(a’)* provides the predominant angle of fiber alignment in a subregion by computing line integrals along parallel-beam projections oriented at discrete rotation angles (*ϴ*) and spaced 1 pixel apart. For a two-dimensional function, *f(a,b)*, *a* and *b* are the horizontal and vertical axes and *a’* and *b’* are the axes of the parallel-beam projections determined by the prescribed rotation angle (*ϴ*). Eqns 3–4 describe the Radon transform:

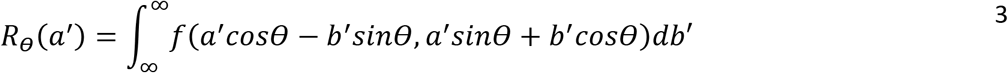

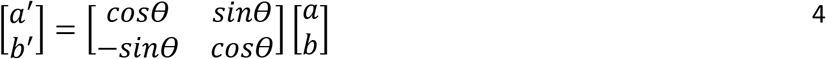

For each image subregion, we computed the Radon transform *R_ϴ_(a’)* while varying rotation angle (*0*°<*ϴ*<*180*°). The subregion Radon transform reaches a unique maximum value at the angle of greatest pixel alignment, and this angle was taken as the subregion predominant collagen fiber direction (*α_i_*) and confined to the first two quadrants such that (*0*°<*α_i_*<*180*°). (**Fig. 2D**).

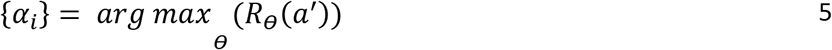

Predominant collagen fiber direction *α_i_* and was calculated for all 4,096 image subregions (**Fig. 2E**).

#### Mean collagen measurements

We utilized built-in MATLAB circular statistics functions^41^ to calculate mean collagen direction and strength of alignment. Subregion collagen directions (*α_i_*) were converted to unit vectors (**r_i_**) and averaged to obtain the image mean resultant vector 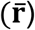.

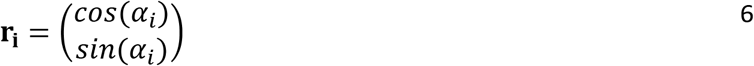

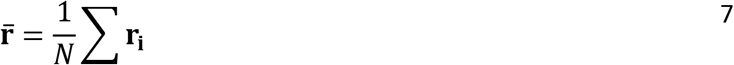

Next, the mean collagen fiber direction (*cfd*) was calculated per image as the acute angle of 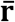 relative to the horizontal axis, such that (*0*°<*cfd*<*90*°). (**Fig. 2H**). The resultant vector length was used to calculate the strength of alignment (*0*<*SA*<*1*, 1=high alignment), capturing the circular spread in local orientations per image.

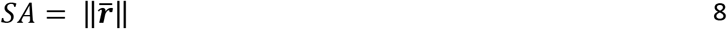

The mean collagen pixel ratio was also determined per image as a measure of the total amount of collagen fibers present.

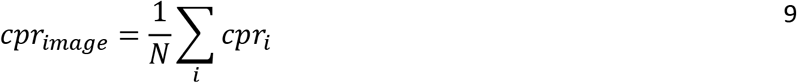

#### Sensitivity and validation

To validate the image processing algorithm and determine the appropriate subregion size, we first used our image processing algorithm with two manually-generated sets of test images of dark lines that approximated collagen fibers. In the first image set, we varied collagen (line) direction over the range *0°<cpd<90*° while holding strength of alignment constant (*SA_known_*=*1*) (**Supp. Fig.1A**). In the second set, we varied strength of alignment over the range *0.92<SA<0.99* with collagen direction constant (*cpd_known_*=*90*°) (**Supp. Fig.1B**). To test the sensitivity of our algorithm to the image subregion size, we also varied the number of image subregions (*2×2<NxN<128×128*) and calculated the error between the known collagen direction and strength of alignment in our test images vs. the values determined by our image processing algorithm. As we increased the number of image subregions, error in collagen direction increased (**Supp. Fig.1C**), with error in strength of alignment minimized at the 64×64 subregion size (**Supp. Fig.1D**). Based on this analysis, we selected the 64×64 subregion size, which yielded a collagen direction error of 0.35° and a strength of alignment error of 0.014.

### In silico finite-element modeling

#### Geometry

We generated finite-element (FE) models in the nonlinear finite element solver, FEBio (Musculoskeletal Research Laboratories, University of Utah, Salt Lake City, UT, USA).^42^ FE models corresponded to the height and width of our cropped SEM images (118μm × 118μm), with a constant thickness (3μm). The FE model was meshed into 64×64 hex8 elements, with each element corresponding to one subregion from our image processing algorithm (**Fig. 3A**).

**Figure 3.**
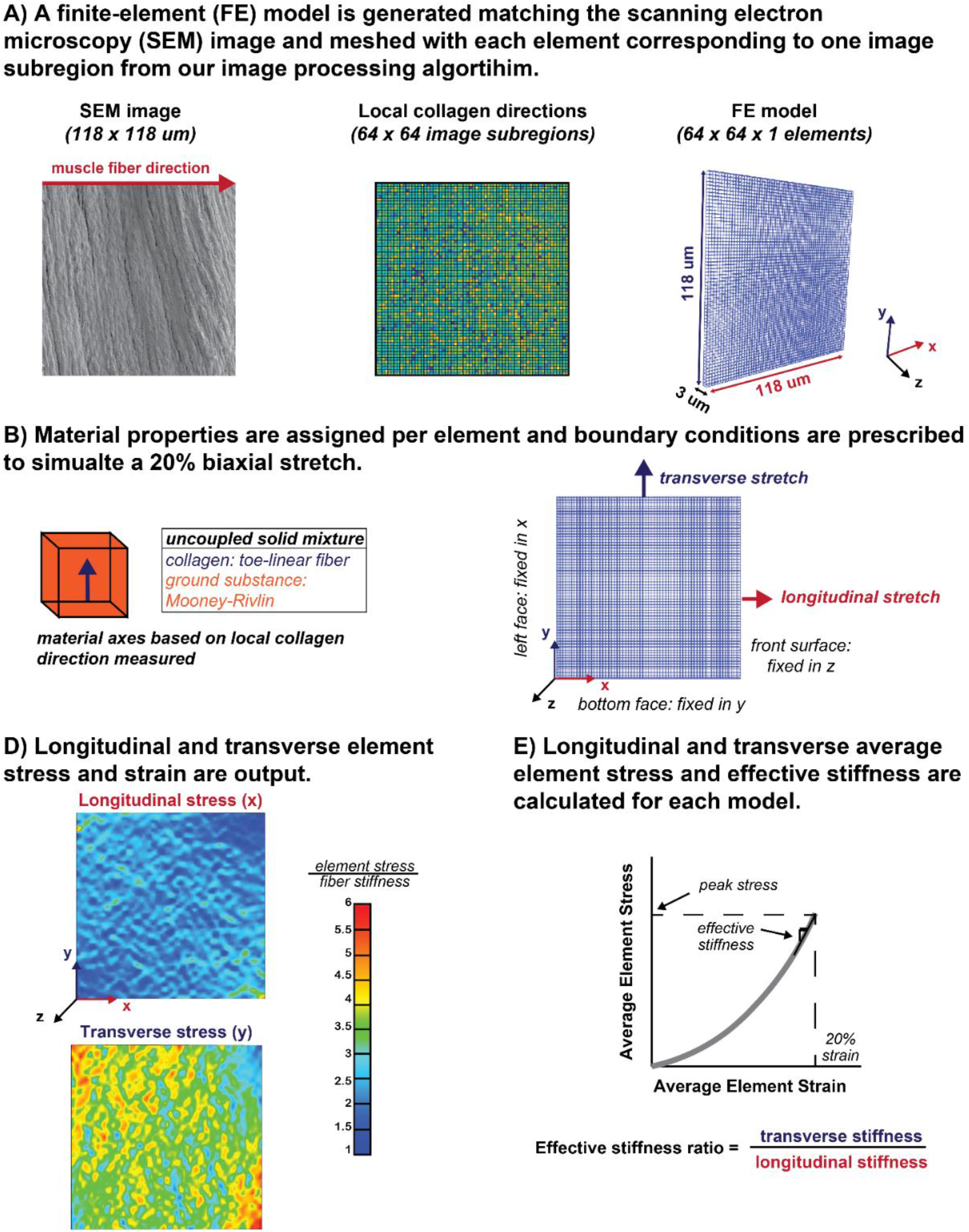
(A) FE models were generated in the nonlinear finite element analysis software suite, FeBio, corresponding to the height and width of our cropped SEM images (118μm × 118μm), with a constant thickness (3μm). The FE model was meshed into 64×64 hex8 elements, with each element corresponding to one image subregion from our image processing algorithm. (B) We assigned a coupled solid mixture constitutive model to the geometry, with material axes assigned per element based on local collagen directions measured in our image analysis. A toe-linear fiber material was used to represent collagen fibers, and a Mooney-Rivlin material was used to represent the remaining ground substance of the ECM. (C) Boundary conditions were assigned to simulate an equibiaxial 20% engineering strain by prescribing displacements to the +y and +x mesh surfaces corresponding to the top and right edge of the SEM image, respectively. The −y surface was fixed in y, the −x surface was fixed in x, and the +z surface was fixed in z. (D) Cauchy stress and Lagrange strain, in the longitudinal (x) and transverse (y) directions, were output for each element and averaged to determine the stress-strain curve for each model. Average element stress was measured at 20% strain in the longitudinal (*S_long_*) and transverse (*S_trans_*) directions. Effective stiffness in the longitudinal (*k_long_*) and transverse (*k_trans_*) directions were measured with a linear fit at the 18, 19, and 20% strain time points. Effective stiffness ratio was then calculated for each model 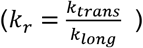.

#### Material law

Connective tissue is often modeled as a composite of collagen fibers embedded in an isotropic “ground matrix”.^43–46^ The “ground matrix” is referred to as a gel-like amorphous substance and contains all non-fibrillar components of the ECM (e.g. proteoglycans, glycosaminoglycans).^12^ Collagen fibers contribute only to the tensile properties of the ECM. Their stress-strain behavior exhibits distinct toe and linear regions under tensile deformation as collagen fibers uncrimp and straighten.^47^ To represent the skeletal muscle ECM, we assigned a coupled solid mixture constitutive model to the geometry. Collagen fibers were modeled as a toe-linear fiber^48^, where the strain energy density is a function of the fiber stretch *λ*. A transition from the toe to linear region occurs at *λ_0_*, where *β* is the power law exponent in the toe region and *E* is the linear fiber modulus. The remaining ground substance of the ECM was modeled as a Mooney-Rivlin material^49^ where *c_1_* and *c_2_* are the Mooney-Rivlin material coefficients and *K* is a bulk modulus-like penalty parameter for the coupled solid mixture. Specific details and constitutive equations of these materials can be found in the FEBio user manual (help.febio.org).

#### Material parameters

The toe region of the collagen fiber stress-strain curve is associated with both the straightening of wavy collagen fibers^50,51^ and collagen fiber realignment,^52^ often modeled with *λ_0_*=*1.06*.^53^ We explicitly modeled collagen fiber straightness by discretizing our image into subregions and measuring local collagen directions. Since collagen fiber straightening is already accounted for in our model, we used a constant stretch ratio of *λ_0_*=*1.01*, such that it only accounts for the contribution of collagen fiber realignment within the image sub-regions. Due to the wide range of values for collagen fiber stiffness and ground matrix stiffness reported in the literature, we varied the ratio of *E* to *c_1_* and conducted a sensitivity analysis described in detail in Supplemental Materials. Based on that analysis, we saw that collagen fiber direction had a greater influence on the effective stiffness ratio as we increased the ratio of *E* to *c_1_*. Therefore, we selected a collagen fiber modulus ten times greater than the ground matrix stiffness (*E*=*10MPa, c_1_*=*1MPa*) so that we were not overestimating the influence of changes in collagen fiber organization. We then selected a bulk modulus to ensure incompressibility (*100*<*K/c_1_*<*10,000*).^46^ The purpose of our FE models was to isolate the effect of ~1μm scale structural changes of collagen fiber organization on ~100 μm scale tissue mechanical behavior. Therefore, the material parameters shown in Table 1 were held constant for all SEM-image based models. Material axes were then assigned per element to reflect the subregion collagen direction measurements (*α_i_*) obtained from our image processing algorithm, with fiber angles in the x-y plane (**Fig. 3B**). The x axis corresponded to the longitudinal (muscle fiber) direction, the y axis corresponded to the transverse direction, and the z axes was orthogonal to the x and y axes.

**Table 1.** Material Parameters for coupled solid mixture material.

| **Toe-linear collagen fiber** |  |  |
| --- | --- | --- |
| E | Fiber modulus in the linear range | 10 MPa |
| $\beta$ | Power-law exponent in the toe region | 3 |
| $\lambda_0$ | Stretch ratio when toe region transitions to the linear region | 1.01 |
| **Mooney-Rivlin ground matrix** |  |  |
| $c_1$ | Mooney-Rivlin $c_1$ parameter | 1 MPa |
| $c_2$ | Mooney-Rivlin $c_2$ parameter | 0 MPa |
| K | Bulk-modulus | 1000 MPa |

#### Boundary conditions

We assigned boundary conditions to simulate an equibiaxial 20% engineering strain (**Fig. 3C**) by prescribing displacements to the +y and +x mesh surfaces corresponding to the top and right edge of the SEM image, respectively. The -y surface was fixed in y, the −x surface was fixed in x, and the +z surface was fixed in z.

#### Model outputs

Cauchy stress and Lagrange strain, in the longitudinal (x) and transverse (y) directions, were output for each element and averaged to determine the stress-strain curve for each model (**Fig.3D**). Average element Cauchy stress was measured at 20% strain in the longitudinal (*s_long_*) and transverse (*s_trans_*) directions. Effective stiffness in the longitudinal (*k_long_*) and transverse (*k_trans_*) directions were measured with a linear fit of the 18, 19, and 20% strain points and effective stiffness ratio was then calculated for each model (*k_r_*= *k_trans_/k_long_*) (**Fig. 3E**). We normalized model outputs of stress and stiffness by collagen fiber modulus (*E*=*10 MPa*) due to uncertainty in estimates for our material parameters.

### Theoretical structure-function relationships from simplified images

To predict the influence of specific parameters of collagen organization on mechanical properties, we simulated changes in each parameter alone in “simplified” images that we manually generated. We used our modeling pipeline to measure the effective stiffness ratio in FE models based on each simplified image. This allowed us to determine “theoretical” structure function relationships and answer questions such as, *“How do collagen fiber straightness and collagen fiber direction influence ECM stiffness ratio independently?”*. By comparing the theoretical relationships with SEM image-based models we then asked questions such as, *“Do changes in collagen fiber straightness or collagen fiber direction measured in SEM images predict changes in effective stiffness ratio predicted in FE models?”,* and *“Do SEM image-based models follow theoretical structure-function relationships?”*.

### Simplified images

First, we created images with varied collagen fiber direction (*0°<cfd<90°*), with collagen fiber straightness constant at 1.0 and then 0.85, since the straightness parameters from SEM images fell within this range (**Supp Fig 2A**). Next, we varied collagen fiber straightness (*0.6<P_s_<1.0*) while holding fiber direction constant at 90° and then 70°, since the collagen fiber directions from SEM images fell within this range (**Supp Fig 2B**). We generated FE models from each simplified image and plotted the effective stiffness ratio vs. the parameter that was varied in the simplified image. This allowed us to fit theoretical curves to the relationships between effective stiffness ratio and collagen fiber direction (**Supp Fig 2C**), as well as between effective stiffness ratio and collagen fiber straightness (**Supp Fig 2D**). We compared the theoretical curves with the results of SEM-image based models to determine whether SEM images of diaphragm muscle ECM followed the theoretical relationship and if functional parameters could be predicted by structural parameters (**Fig. 8**). The image processing and modeling code will be made publicly available upon publication.

### Statistical analysis

A two-way analysis of variance (ANOVA) with age (3, 6, 12 months) and group (*mdx* vs WT) as factors was performed for the following measurements: (1) collagen pixel ratio, (2) collagen fiber direction relative to muscle fiber direction, (3) collagen fiber straightness parameter, (4) strength of alignment, (5) effective stiffness (longitudinal and transverse), (6) average element stress (longitudinal and transverse), (7) effective stiffness ratio. Assumptions of random sampling, equal variance, and normality of residuals were confirmed with qq plots and distribution plots. When applicable, Tukey HSD post-hoc comparison was performed to determine which groups were significantly different. Alpha was set at 0.05 for all tests. Exponential and power law curves were fit to our simulated models and experimental data with nonlinear regression (MATLAB (R2018b) and Curve Fitting Toolbox 3.5.8).^54^

## 3. RESULTS

### Collagen fibers are straighter and more highly aligned in *mdx* and older WT mice but retain a transverse orientation relative to muscle fibers

Changes in collagen fiber organization can be detected visually in the SEM images (**Fig.4**). Collagen fiber straightness was significantly greater in *mdx* over WT at 3 months (*mdx*=0.976±0.0108, WT=0.887±0.0309, p=3.3e-6) and 6 months (*mdx*=0.942±0.0182, WT=0.881±0.0163, p=1.0e-3). Collagen fiber straightness was also significantly greater in 12-month-old WT (0.931±0.0289) over 3-month-old WT (0.887±0.0309), (p=0.027), as well as 12-month-old WT (0.931±0.0289) over 6-month-old WT (0.881±0.0163), (p=0.0090) (**Fig. 5A**). Collagen fiber strength of alignment was significantly greater in *mdx* over WT groups at 3 months (*mdx*=0.876±0.0333, WT=0.759±0.0416, p=3.0e-5) and 6 months (*mdx*=0.840±0.0315, WT=0.759±0.0368, p=4.5e-3) (**Fig. 5B**). Collagen pixel ratio ranged from 0.47-0.61, with no significant differences between age or disease groups (**Fig. 5C**). Collagen fiber direction relative to muscle fiber direction ranged from 70-90°, with no significant differences between age or disease groups (**Fig. 5D**).

**Figure 4:**
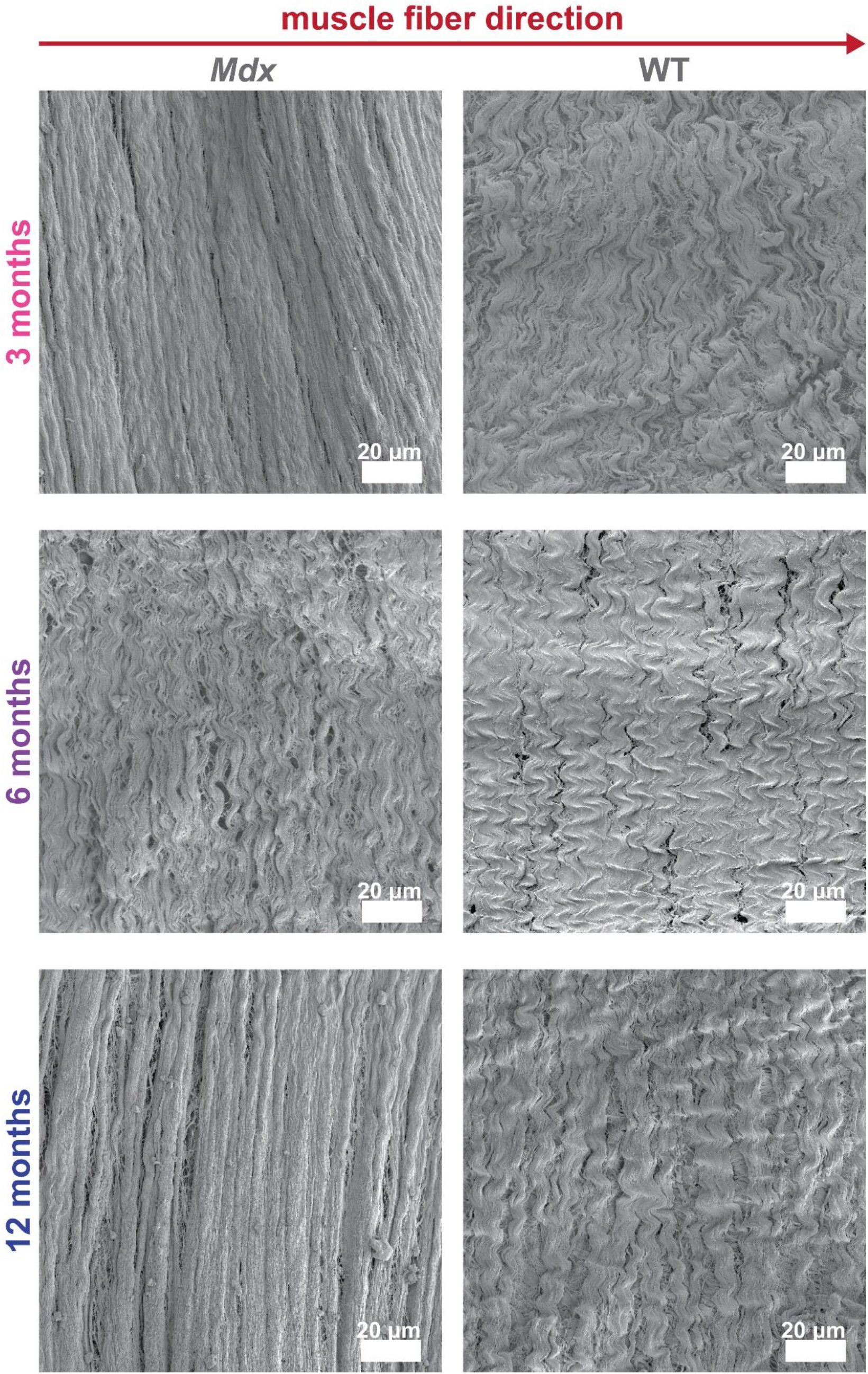
Diaphragm muscle samples were dissected from the costal region and collagen structure was isolated after muscle fiber digestion. Samples were imaged with Scanning Electron Microscopy (Zeiss Sigma VP HD field SEM) and images collected at 1kX were used to visualize collagen fiber organization. We assume a stress free, fully relaxed configuration for our samples from 3, 6, and 12-month old *Mdx* and WT (n=6) mice.

**Figure 5:**
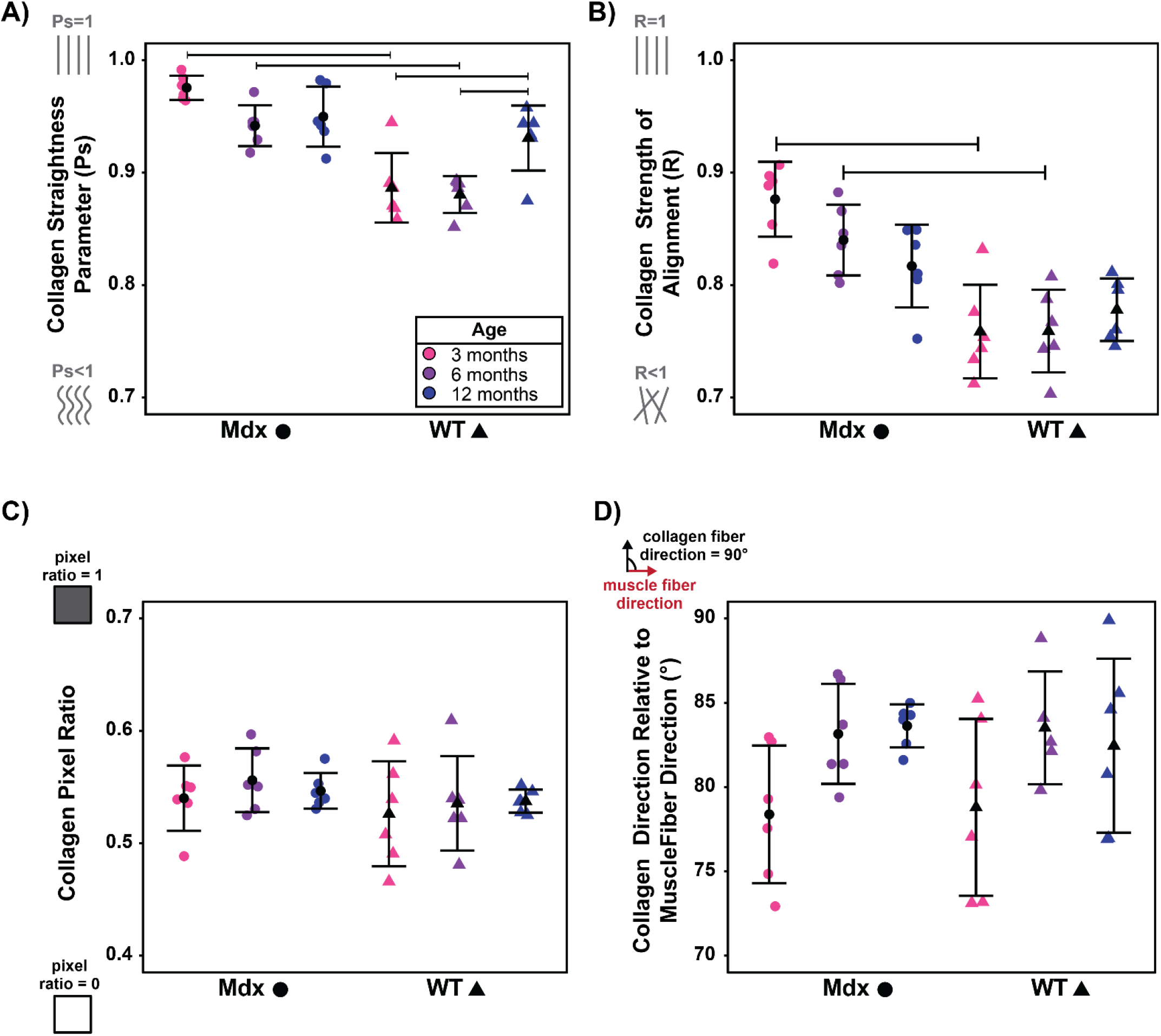
(A) Collagen fiber straightness (*P_s_*) measured for three fibers per image and averaged. (B) Collagen strength of alignment (*SA*) used to measure the circular spread in local orientations per image (*0*<*SA*<*1*, 1=high alignment). (C) Collagen pixel ratio, quantified per image as the number of collagen pixels divided by the number of total pixels. (D) Mean collagen orientation relative to the muscle fiber direction, measured as the average of local directions measured per image window. Significance between groups shown with bars, where p<0.05 and n=6 mice per group.

### The mechanical models predict that longitudinal average stress and effective stiffness are greater in WT mice, and transverse average stress and effective stiffness are greater in *mdx* mice

Variations in collagen fiber organization measured in SEM images were reflected qualitatively in the element stresses in the FE models. (**Fig.6**). Longitudinal average stress was significantly greater in WT over *mdx* at 3 months (*mdx*=0.233±0.0116, WT=0.262±0.0107, p=2.2e-4) and 6 months (*mdx*=0.239±0.00762, WT=0.261±0.00948, p=4.4e-3) (**Fig. 7A**). Transverse average stress was significantly greater in *mdx* over WT at 3 months (*mdx*=0.347±0.0161, WT=0.311±0.0135, p=4.0e-4) and 6 months (mdx=0.340±0.0105, WT=0.310±0.0118, p=3.8e-3) (**Fig. 7B**). Longitudinal effective stiffness was significantly greater in WT over *mdx* at 3 months (mdx=0.900±0.0541, WT=1.037±0.0510, p=1.9e-4) and 6 months (*mdx*=0.928±0.0363, WT=1.035±0.0456, p=4.1e-3) (**Fig. 7C**). Transverse effective stiffness was significantly greater in *mdx* over WT at 3 months (*mdx*=1.447±0.0809, WT=1.267±0.0675, p=4.3 e-4) and 6 months (*mdx*=1.415±0.0532, WT=1.265±0.0604, p=4.0 e-3) (**Fig. 7D**).

**Figure 6:**
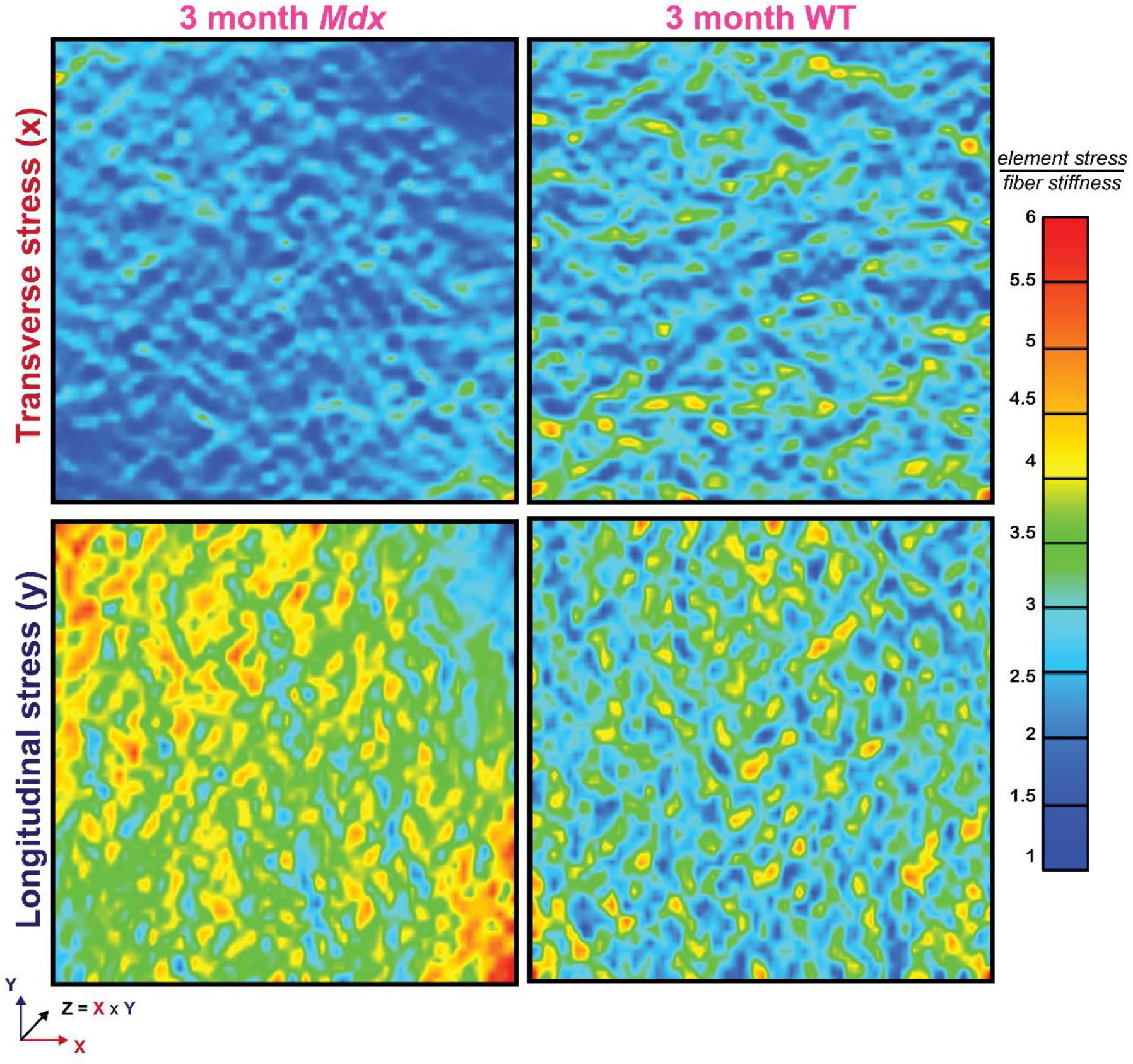
Finite-element models based on images of 3-month-old *mdx* and WT mice (seen in Figure 4). Element stress normalized by fiber stiffness is plotted in the transverse (top) and longitudinal (bottom) directions.

**Figure 7:**
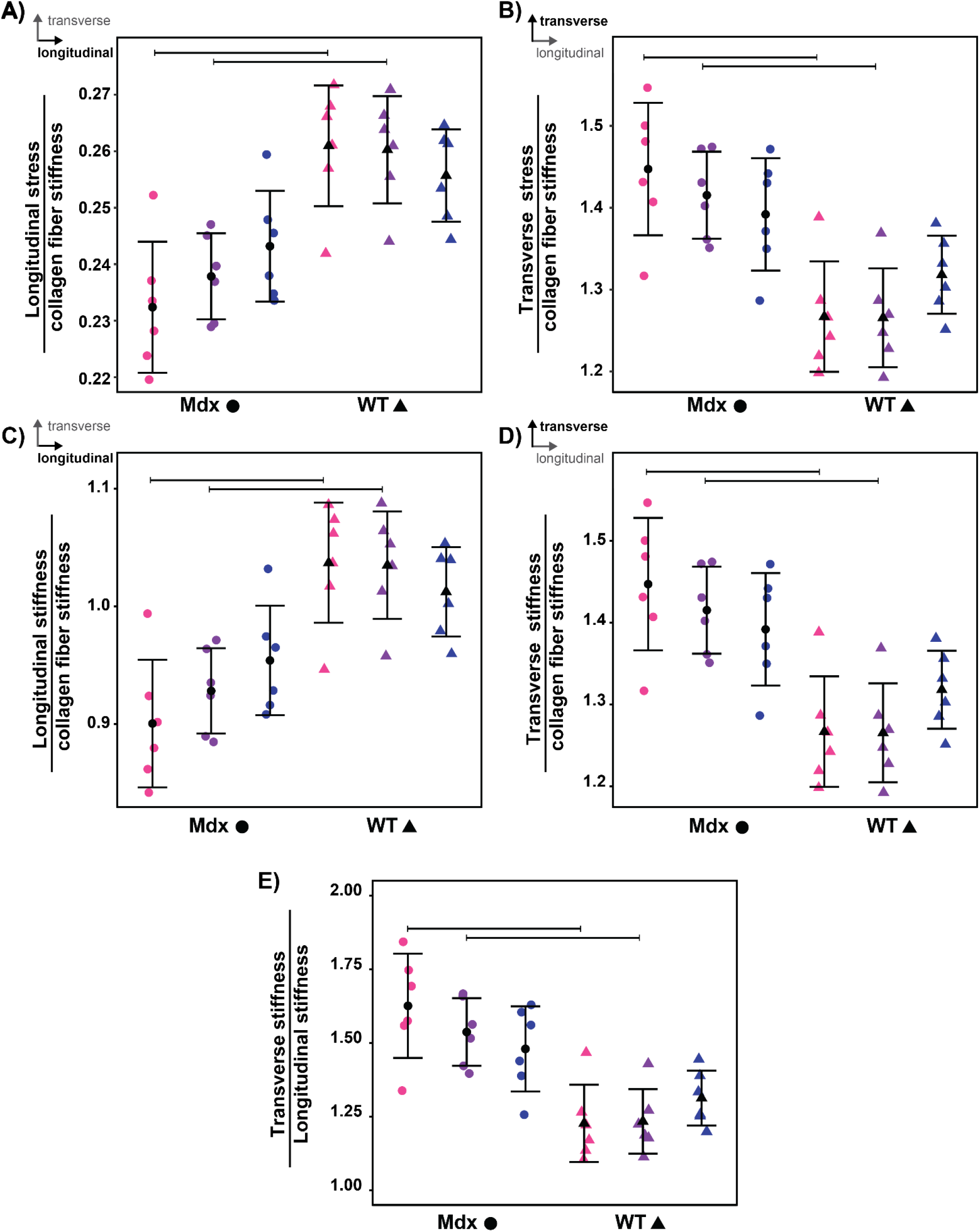
(A) Longitudinal average element stress normalized by fiber stiffness. (B) Transverse average element stress normalized by fiber stiffness. (C) Longitudinal effective stiffness at 20% strain, normalized by fiber stiffness. (D) Transverse effective stiffness at 20% strain, normalized by fiber stiffness. (E) Stiffness ratio quantified as the transverse stiffness divided by longitudinal stiffness. Significance between groups shown with bars, where p<0.05 and n=6 mice per group.

### The ratio of transverse to longitudinal stiffness was greater in *mdx* mice and follows a theoretical power law relationship with collagen fiber straightness

For all SEM-image based models, the effective stiffness ratio was greater than 1, indicating greater stiffness in the direction transverse to the muscle fibers. The effective stiffness ratio was significantly greater in *mdx* over WT at 3 months (*mdx*=1.626±0.177, WT=1.227±0.131, p=1.5e-4) and 6 months (*mdx*=1.537±0.115, WT=1.234±0.109, p=4.6e-3) (**Fig. 7E**). FE models based on simplified images reveal theoretical power law relationships between collagen fiber straightness and effective stiffness ratio and exponential relationships between collagen direction and stiffness ratio (**Fig. 8**). SEM-image based models follow the theoretical power law relationship between collagen straightness and stiffness ratio (*k_r_*=*0.8224xPs^9.653^* + *0.9744, R^2^*=*0.7092*), with predicted values falling below both simulated curves (**Fig. 8A**). SEM-image based models do not fit to a theoretical exponential relationship between collagen direction and stiffness ratio, with data points falling between the two theoretical curves (**Fig. 8B**).

**Figure 8:**
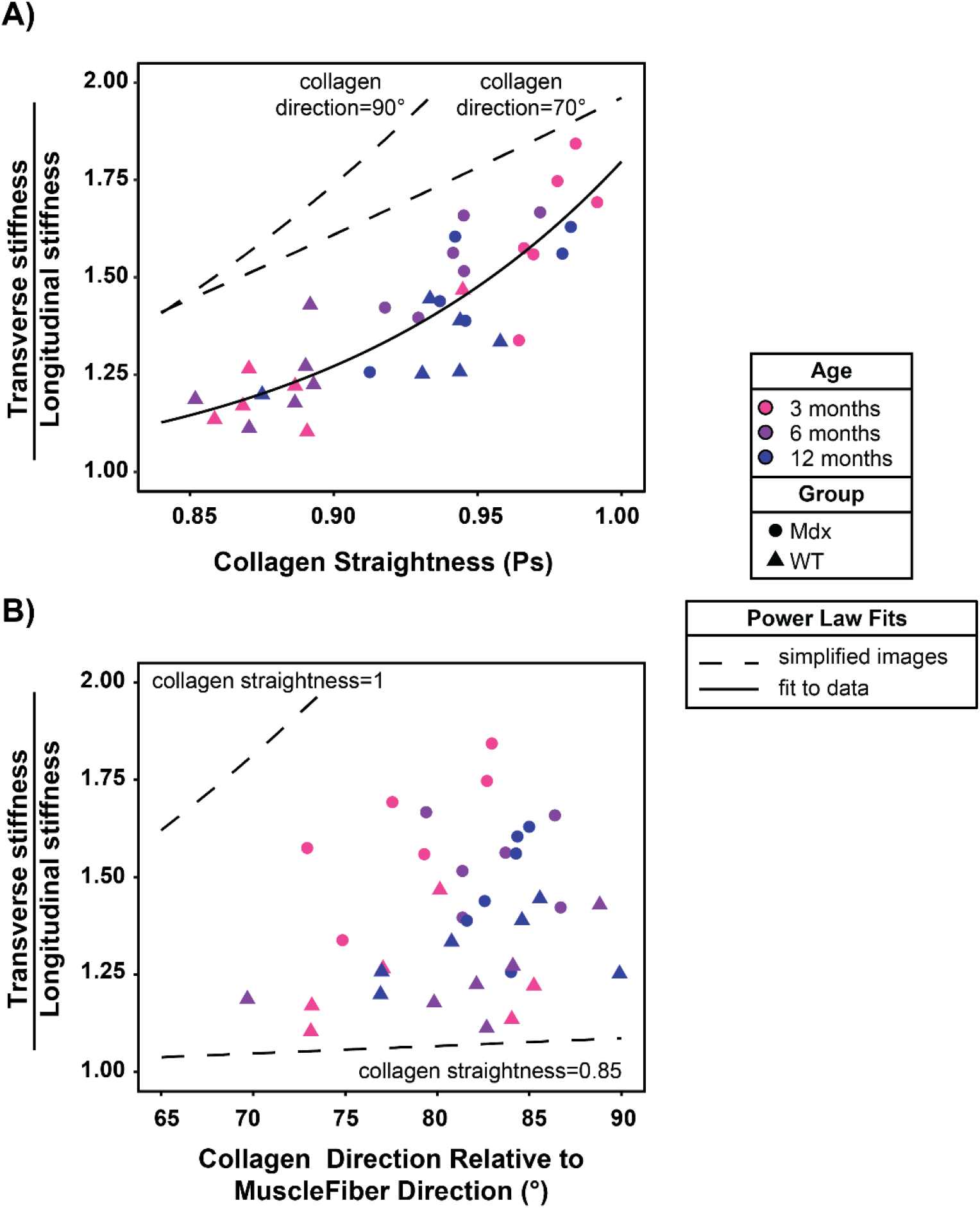
(A) Relationship between collagen straightness and stiffness ratio from diaphragm muscle samples, as well as simplified images where we varied collagen fiber straightness with collagen direction constant at 70 or 90°. Power law curves fit to simplified images (*R^2^*>*0.99*) and SEM image-based model (*kr*=*0.8224xPs9.653* + *0.9744, R^2^*=*0.7092*). (B) Relationship between collagen fiber direction and stiffness ratio from diaphragm muscle samples, as well as simplified images where we varied collagen direction with collagen fiber straightness constant at 1 or 0.85. Exponential curve fit to simplified images (*R^2^*>*0.97*).

## 4. DISCUSSION

In this study we tested the hypotheses that collagen structure within the ECM is altered in DMD and that these changes have implications of the mechanical properties of the ECM. We first visualized collagen structure with SEM images and then developed an analysis framework to quantify collagen organization and explore the influence of our measurements on ECM mechanics (**Fig.1–3**). The image analysis reveals that collagen fibers within the diaphragm muscle epimysium are oriented transversely, with increased collagen fiber straightness and alignment with age and disease (**Fig.5**). From the SEM image-based mechanical models, we predict that transverse effective stiffness and average stress are also increased with age and disease. Additionally, both healthy and diseased models reveal an increase in transverse stiffness relative to longitudinal stiffness, with the ratio of transverse to longitudinal stiffness increased with disease (**Fig.7**). From the models based on simplified images, we predict a theoretical power law relationship between collagen fiber straightness and stiffness ratio (**Fig.8A**) and a theoretical exponential relationship between collagen direction and stiffness ratio (**Fig.8B**). By comparing with the SEM image-based models, we see that collagen straightness is a significant predictor of stiffness ratio (**Fig. 8A**) while collagen direction is not (**Fig. 8B**).

### Our findings implicate changes in ECM structure and mechanics on the mechanical properties of diaphragm muscle

As DMD progresses in patients, quantitative measures such as respiratory Forced Vital Capacity show a decline in pulmonary function with age.^55^ The respiratory muscles are responsible for mechanical regulation of breathing, and the diaphragm is the principal inspiratory muscle separating the thoracic and abdominal regions. As the diaphragm muscle contracts and shifts downwards, air is drawn into the lungs during inspiration. As the diaphragm muscle relaxes, air is pushed out of lungs during expiration. The contribution of the abdomen to the tidal lung volume is used as a marker of diaphragm muscle weakening, which decreases with age in DMD patients.^56^ Diaphragm muscle excursion, the vertical displacement between the diaphragm position at inspiration and expiration, is another marker of diaphragm dysfunction and also decreases with age in DMD patients.^10^ To understand how tissue level properties lead to a changes in muscle function, animal models such as the *mdx* mouse allow us to measure changes in mechanical properties with disease. Mechanical properties of diaphragm muscle are often measured from uniaxial strip tests and show a decrease in elasticity and force in *mdx* diaphragm.^18,57^ However, unlike most skeletal muscles, the diaphragm sustains biaxial loads *in vivo* and exhibits nonuniform and anisotropic behavior.^58–60^ From uniaxial tests of rat diaphragm muscle, Boriek et al. report that extensibility of diaphragm tissue is decreased when muscle is loaded uniaxially transverse to muscle fibers, than when loaded uniaxially along muscle fibers.^59^ From biaxial tests of canine diaphragm muscle, Boriek et al. report an increase in transverse relative to longitudinal passive stiffness.^60^ In this study, models simulated a biaxial test and predicted that transverse stiffness was greater than longitudinal stiffness for both healthy and *mdx* mice, similar to Boriek et al.’s findings in canine diaphragm muscle tissue.^60^

### The methods presented here offer a novel framework to explicitly model the influence of ECM structure on mechanical properties

Gao et al. developed a constitutive model^61^ to explain age related stiffening of epimysium from rat tibialis anterior muscle where no differences in collagen structure were detected in SEM images.^62^ In their model, they represented collagen fibers with unit cells and assigned unit cell angle from collagen fiber distributions that were measured experimentally^34^ but were not specific to the epimysium. They found that increased tissue level stiffness was explained by increased stiffness of collagen fibers and ground matrix rather than the geometry or distribution of collagen fibers. In the modeling pipeline presented here, we explicitly modeled collagen organization by assigning fiber directions in each finite element, allowing for a framework that can be easily translated. In the SEM images, we found an increase in collagen fiber straightness in older healthy mice relative to younger healthy mice (**Fig. 5A**), but this difference was not reflected in the model predictions (**Fig.7**). In both images and models, we did not find any differences between diseased and healthy groups at 12 months, suggesting that the ECM of older healthy mice resembles that of diseased mice. A benefit of our modeling pipeline is that we can isolate structural parameters using simplified images and predict structure-function relationships from our FE models. The models based on SEM images follow the theoretical power law relationship between collagen straightness and stiffness ratio, but the experimental data falls below both predicted curves (**Fig. 8A**). This suggests that there is isotropy beyond changes in collagen fiber straightness accounting for an increase in stiffness ratio in the diseased groups. The models based on SEM images did not follow the theoretical exponential relationship between collagen straightness and stiffness ratio, but data points fell between the predicted curves (**Fig. 8B**). This finding suggests that although collagen fiber direction influences the stiffness ratio, it does not explain differences in model predictions for *mdx* and WT mice. Taken together, these findings suggest that although collagen fiber direction determines the direction in which the ECM will be stiffer in tension, collagen fiber straightness regulates the ratio of transverse to longitudinal stiffness.

There are some limitations of this approach that should be mentioned. Enzymatic and detergent digestions such as the sodium hydroxide protocol used in this study have been shown to alter the mechanical properties of the ECM.^27^ While this is what motivated us to use modeling to explore the mechanical implications of the ECM, changes in the structure of collagen fibers may have been influenced by removing all other ECM components. Since all samples underwent the same protocol, we assume that any influence was constant between groups. In the FE models, we grouped all collagen subtypes in one material and did not account for changes in other ECM components or crosslinking of collagen fibers. For this reason, it is important to acknowledge that the results presented in this work focus on the effects of collagen fiber organization alone. Additionally, we did not measure mechanical properties of our decellularized samples directly. While simplistic, this allowed us to focus specifically on relating structural parameters at the collagen fiber level to bulk level tissue properties. Thus, the modeling results presented here are theoretical in nature and provide interesting hypotheses for future experiments to examine changes in passive mechanics of the dystrophic diaphragm.

### The skeletal muscle ECM is a complex three-dimensional scaffold that is organized uniquely across muscle groups

Skeletal muscle fibrosis is often characterized with images of muscle cross-sections, capturing the perimysium, surrounding muscle fascicles, and endomysium, surrounding muscle fibers.^34,63,64^ A honeycomb structure is seen, with a build-up of collagen often quantified by collagen area fraction from these cross-sectional images. The structure of the endomysium is similar across skeletal muscles,^65^ while differences in perimysium are reported across muscle groups and with disease. Borg et. al report that perimysium from diaphragm muscle is less developed than other skeletal muscle groups where large bundles of collagen fibers are seen, arranged both parallel and circumferential to muscle fibers.^65^ An increase in the number of such “perimysial collagen cables” has been reported with fibrosis, ^66^ but their prevalence in diaphragm muscle fibrosis remains unknown. In our study, we collected images of the diaphragm muscle epimysium, the outermost layer of the ECM surrounding skeletal muscle. Differences in the structure of the epimysium are also reported across skeletal muscles. In long strap-like muscle, a cross-ply arrangement of wavy collagen fibers oriented approximately 55° to the muscle fiber direction is reported, and in pennate muscle a dense layer of collagen fibers aligned parallel to muscle fibers is reported.^63^ In Gao et al.’s study of the epimysium from young and old rat tibialis anterior muscles, they report that the outer layer consists of wavy collagen fibers that were highly aligned in a “predominant direction” but do not report the direction relative to muscle fiber direction and do not quantify collagen alignment or straightness.^62^ Our SEM images show a similar arrangement to Gao at al. in the diaphragm muscle epimysium, but with collagen fibers oriented transverse to muscle fibers. We hypothesize that this arrangement of collagen fibers is unique to the diaphragm muscle and highlights the need to study the arrangement of collagen fibers in each muscle before we can determine how changes during fibrosis affect its mechanical properties.

### The skeletal muscle ECM is essential for transmitting forces from muscle fibers

Muscle fibers can transmit force both longitudinally along the muscle fiber axis, and laterally to adjacent muscle fibers.^67–69^ Lateral force transmission occurs through physical linkages between the actin cytoskeleton and ECM at the muscle fiber membrane.^70^ This notion is supported by the ability of the endomysium to transmit forces between intrafascicularly terminating muscle fibers through shear^69^ and the physical continuity of the perimysium from muscle to tendon.^71^ Huijing et al. describe “epimuscular myofascial force transmission” as the force transmitted between muscle and its surroundings through the epimsuyim.^72^ This idea is supported through experiments where force transmission still occurs after tendonotomy^73,74^ and differences in proximal and distal force are measured when the muscle-tendon complex length is held constant.^75^ Such experiments provide strong evidence of force transmission through pathways other than the myotendinous junctions and highlight the role of the ECM. Damage to connective tissue is shown to hinder force transmission and we must consider how fibrosis affects the ability of the ECM to transmit forces.^75^ In *mdx* mice, where the muscle fiber membrane is weakened due to the lack of dystrophin, lateral force transmission is severely impaired.^68^ The diaphragm muscle has a complex architecture with a majority of intrafascicularly terminating muscle fibers,^76^ suggesting that the ECM is especially critical for lateral force transmission. The SEM images in this study revealed that collagen fibers are oriented transverse to the muscle fibers, suggesting that they may serve as a direct pathway for transmitting forces laterally between muscle fibers. Thus, changes in the alignment or straightness of these transversely oriented collagen fibers may influence the ability of the ECM to transmit lateral forces between neighboring muscle fibers of from muscle fibers to tendon.

### We must consider the role of collagen organization on respiratory insufficiency in DMD

To hypothesize the implications of our findings on respiration we must first consider the unique architecture of the diaphragm muscle. The diaphragm is a dome shaped, “sheet-like” muscle, with a central tendon connecting to the costal and crural muscle domains.^77^ In our study, we collected samples from the costal region. In this region, the muscle fibers are arranged radially from the central tendon to insertion at the rib cage. Therefore, if we consider our results at the whole-muscle level we expect that collagen fibers are arranged circumferentially to maintain the transverse orientation we measured at high magnifications. Collagen fibers are responsible for generating force when stretched in tension, implicating that the epimysium limits circumferential expansion of the diaphragm muscle. This suggests that the epimysium may regulate the ability for the diaphragm muscle to return to its fully relaxed configuration during expiration. To better understand the mechanical role of the epimysium on respiratory insufficiency we must characterize the changes in biaxial properties of diaphragm muscle with fibrosis and develop methods to quantify *in vivo* motion of the diaphragm during respiration.

### Future work should consider the unique structure and mechanics of fibrotic tissue in therapies for DMD

Despite progress in recent years, DMD remains a fatal condition, with fibrosis a key contributor to muscle dysfunction and hypothesized to decrease the effectiveness of therapeutics. Anti-fibrotic therapies target inflammatory pathways such as TGF-β, but their effectiveness is measured by decreasing levels of collagen expression, without accounting for changes in ECM structure or mechancis.^78,79^ The importance of mechanical and structural properties of the ECM is well documented in the literature, with ECM stiffness and alignment key regulators of cellular behaviors involved in fibrosis, such as fibroblast alignment, migration, and proliferation.^22,80^ Our study reveals changes in both ECM structure and mechanics in fibrosis, highlighting the need to study the role of therapeutics on collagen organization. Further, we must account for differences in collagen organization between tissue systems, especially as we aim to alleviate the deleterious impacts of fibrosis in DMD in the diaphragm muscle.

## SUPPLEMENTAL INFORMATION

**Supplemental Figure 1:**
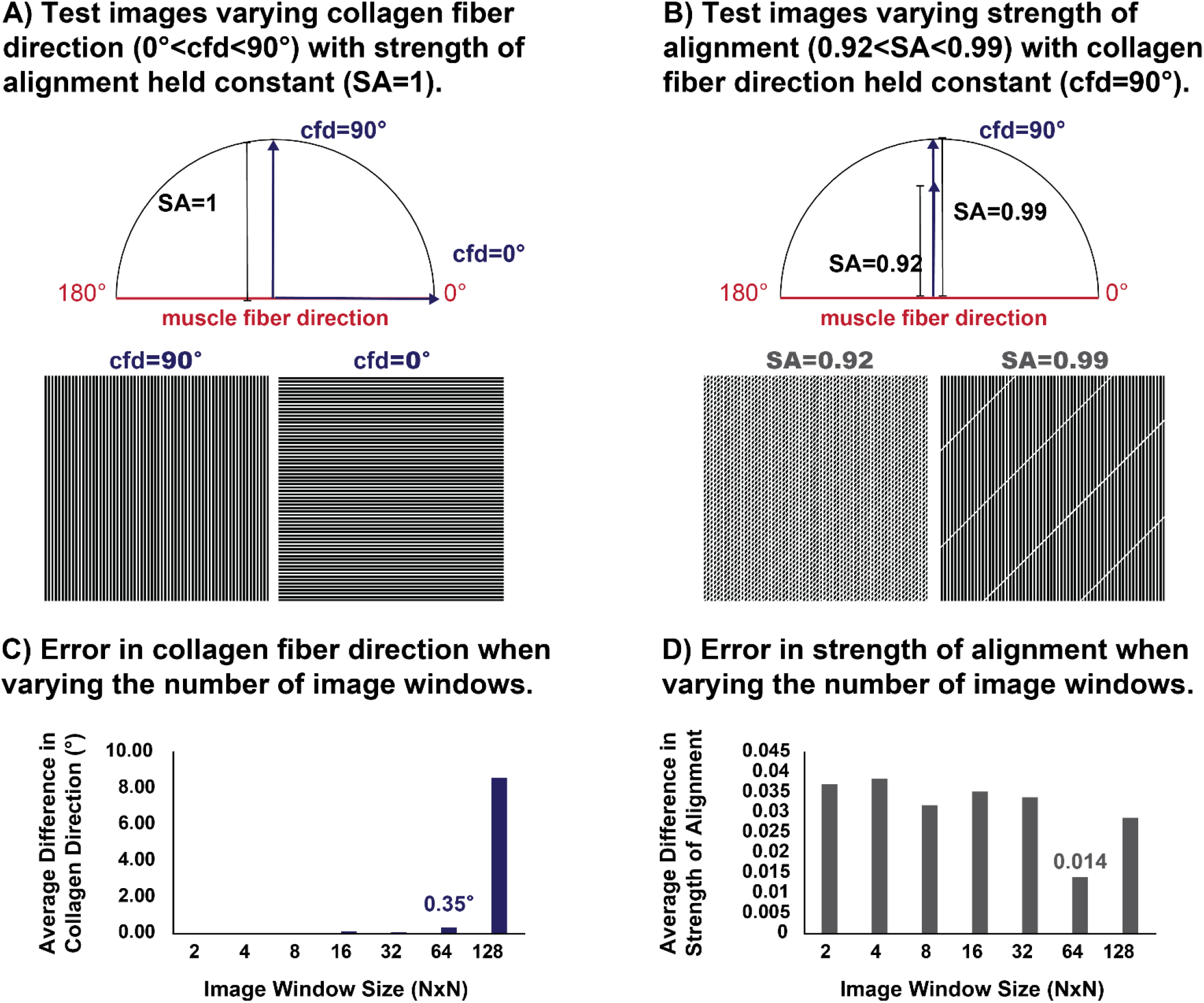
To validate the image processing algorithm and determine the appropriate subregion size, we first used our image processing algorithm with two manually-generated sets of test images of dark lines that approximated collagen fibers. (A) In the first image set, we varied collagen (line) direction over the range *0°<cpd<90°* while holding strength of alignment constant (*SA_known_*=*1*). (B) In the second set, we varied strength of alignment over the range *0.92<SA<0.99* with collagen direction constant (*cpd_known_*=*90*°). To test the sensitivity of our algorithm to the image subregion size, we also varied the number of image subregions (*2×2<NxN<128×128*) and calculated the error between the known collagen direction and strength of alignment in our test images vs. the values determined by our image processing algorithm. (C) As we increased the number of image subregions, error in collagen direction increased. (D) Error in strength of alignment was minimized at the 64×64 subregion size. Based on this analysis, we selected the 64×64 subregion size, which yielded a collagen direction error of 0.35° and a strength of alignment error of 0.014.

**Supplemental Figure 2:**
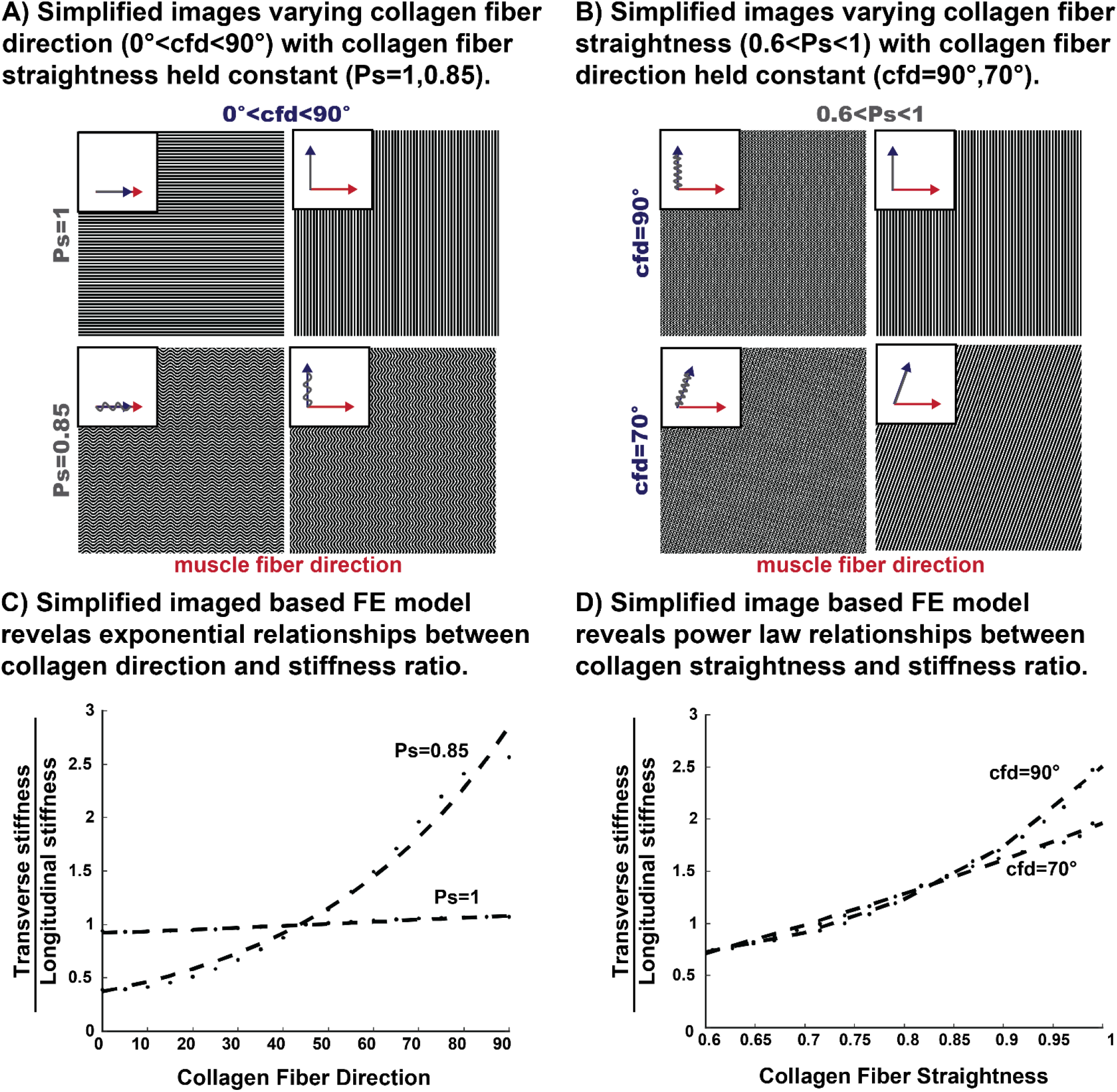
A) First, we varied collagen fiber direction (*0°<cfd<90°*), with collagen fiber straightness constant at 1.0 and then 0.85, since the straightness parameters from SEM images fell within this range. B) Next, we varied collagen fiber straightness (*0.6<P_s_<1.0*) while holding fiber direction constant at 90° and then 70°, since the collagen fiber directions from SEM images fell within this range. C) We generated FE models from each simplified image in A and plotted the effective stiffness ratio vs. collagen fiber direction for each model. We fit exponential relationships for each curve (*R^2^*>*0.99*). D) We generated FE models from each simplified image in B and plotted the effective stiffness ratio vs. collagen fiber straightness for each model. We fit power law relationships for each curve (*R^2^*>*0.97*).

### Supplemental Section 3

A wide range of values for collagen fiber stiffness are reported in the literature, varying from 300 MPa-12 GPa^81–83^ and are dependent on the mechanical testing conditions. Estimates for ground matrix stiffness also have a wide range, varying from 0.01-1 MPa.^61,84^ These material parameters are often fit to experimental data. In this study we were specifically interested in the influence of collagen fiber organization on tissue level properties alone. Therefore, we varied collagen fiber stiffness relative to ground matrix stiffness to determine the sensitivity of our model outputs to the ratio of *E/c_1_*. We held ground matrix stiffness constant (*c1=1MPa*) and varied the collagen fiber modulus relative to the ground matrix stiffness (*10<E/c_1_<100*). We generated FE models from test images of where we varied collagen direction over the range *0°<α<90°* (**Supp Fig.1A**) and compared the model output of effective stiffness ratio. As *E/c_1_* increased, we saw that collagen fiber direction had a greater influence on the effective stiffness ratio. Therefore, we selected *E/c_1_=10* so that we were not overestimating the influence of changes in collagen fiber organization.

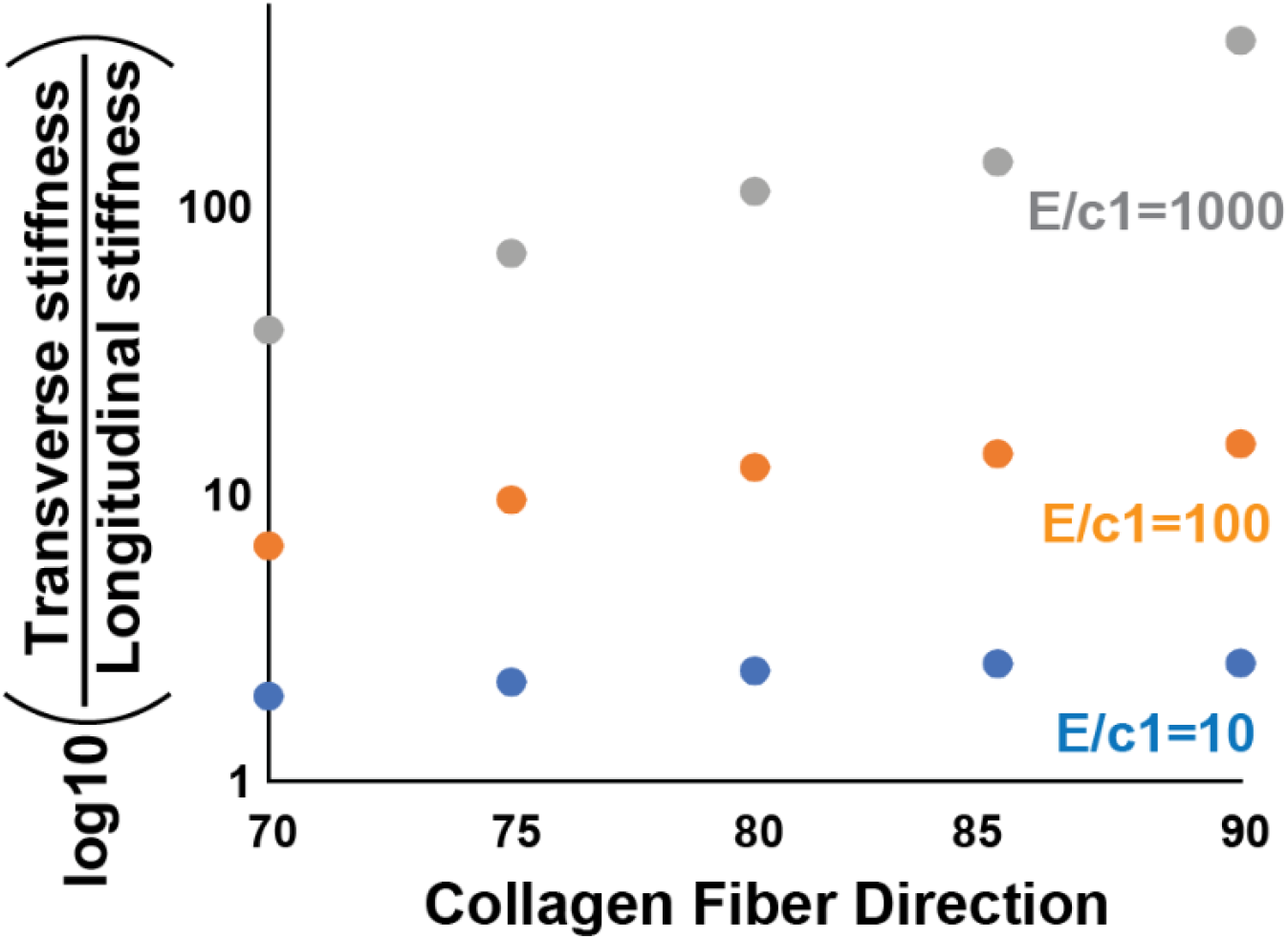

